# Temporary mate removal during incubation leads to variable compensation in a biparental shorebird

**DOI:** 10.1101/117036

**Authors:** Martin Bulla, Mihai Valcu, Anne L. Rutten, Bart Kempenaers

## Abstract

Biparental care for offspring requires cooperation, but it is also a potential source of conflict, since one parent may care less at the expense of the other. How, then, do parents respond to the reduction of their partner’s care? Theoretical models predict that parents that feed offspring should partially compensate for the reduced care of their partner. However, for incubating birds partial compensation is unlikely the optimal strategy, because the entire brood can fail with reduced care. Although biparental incubation dominates in non-passerine birds, short-term manipulations of parental care and evaluations of individual differences in the response, both crucial to our understanding of parental cooperation, are scarce. Here, we describe the response of semipalmated sandpiper (*Calidris pusilla*) parents to a 24-hour removal of their partner during the incubation period, explore factors that can explain individual variation in the response, and describe how incubation rhythms changed after the removed parent returned. On average, the parents compensated partially for the absence of their partner’s care (59%; 95%CI: 49-70%, *N* = 25 individuals). The level of compensation did not differ between the sexes. However, individual responses varied from no to full compensation (2-101%). In seven nests where the removed parent never returned, the widowed parent attended the nest for 0-10 days (median: 4 days). In contrast to patterns observed in undisturbed nests or in uniparental species, nest attendance during compensation tended to be higher during the warmer part of the day. Although the level of compensation was not related to the before-experimental share of incubation, more responsive parents (that left the nest earlier upon human approach) compensated more. The quality of incubation in the after-experimental period was lower than usual, but improved quickly over time. Our findings suggest that full compensation might be limited by energetic constraints or by variation in responsiveness to the absence of the partner. Nevertheless, semipalmated sandpiper parents are able to adjust their subsequent incubation behaviour to take full responsibility for the nest when widowed. Because (nearly) full compensation was the most common response, we speculate that all individuals attempt full compensation, but that some fail because their energy stores get depleted, or because they are less responsive to the absence of their partner.

## INTRODUCTION

Biparental care can be seen as a complex social behaviour where females and males cooperate in the rearing of their offspring. Whereas both parents gain from parental care provided by either of the parents, each parent only pays the costs of its own care. Consequently, each parent would have higher overall reproductive success if the other parent provided a larger share of the care (Trivers 1972; Lessells 2012). How do parents achieve cooperation in face of this conflict?

Established theoretical models predict that parents should partially compensate for a reduction in their partner’s care when an increase in parental care increases breeding success, but with diminishing returns (Houston & Davies 1985; McNamara, Gasson & Houston 1999; McNamara *et al.* 2003; reviewed by Lessells 2012). Partial compensation indeed seems to prevail when considering chick feeding in passerine birds (Harrison et al. 2009), a context for which the models were initially developed. However, partial compensation is unlikely when breeding attempts fail due to a small decrease in parental care, i.e. when the return of investment is zero unless a threshold amount of care is delivered. Such a situation is typical for biparental incubation of eggs in birds, especially in species where the presence of both parents is essential for successful reproduction and in extreme environments where unattended eggs are at high risk of predation (e.g. in gull or frigatebird colonies; Dearborn 2001; Jones, Ruxton & Monaghan 2002), overheating (e.g. in deserts; AlRashidi *et al.* 2011) or severe cooling (e.g. in the Arctic or Antarctica; Gauthier-Clerc *et al.* 2001; Bulla *et al.* 2014). Here, a theoretical model predicts that parents should compensate fully or not at all (Jones, Ruxton & Monaghan 2002).

An experiment to test this prediction of full or no compensation needs to be short term, because parents of biparental species incubate their clutch nearly continuously and hence they might not be able to compensate for the entire remaining incubation period (Deeming 2002). Furthermore, because some individuals may compensate fully while others not at all, evaluating the results of such an experiment at the population level would blur this dichotomy by suggesting partial compensation where none of the individuals partially compensate. Thus, the results need to be evaluated at the individual level.

Biparental incubation of eggs prevails in 50% of avian families (and in 80% of non-passerine ones; Deeming 2002) and many experimental studies have targeted cooperation during biparental incubation. However, these studies are dominated by two approaches with long-term and irreversible effects for the focal breeding attempt. The first approach is to completely remove one parent, creating a situation of no care, reflecting permanent nest desertion (Burley 1980; Erckmann 1981; Bowman & Bird 1987; Brunton 1988; Duckworth 1992; Pinxten, Eens & Verheyen 1995). The second approach is to handicap one parent, creating a situation of reduced care, for example by experimentally increasing plasma testosterone levels in males (De Ridder, Pinxten & Eens 2000; Alonso-Alvarez 2001; McDonald, Buttemer & Astheimer 2001; Schwagmeyer, Schwabl & Mock 2005) or by attaching weights to one of the parents (Wiebe 2010). In contrast, short-term, reversible manipulations of female and male incubation effort (e.g. by supplemental feeding or temporary removal of one parent) with evaluation of between-individual differences in response to this manipulation are scarce (Gibbon, Morrell & Silver 1984; Kosztolányi, Székely & Cuthill 2003; Kosztolanyi, Cuthill & Szekely 2009). The advantage of studying cooperation and solutions to conflict with a short-term, reversible manipulation is not only that it allows testing for full or no compensation, but also that it mimics natural short-term deficiencies in the partner’s care. Such manipulations will show an individual’s immediate response to a sudden reduction in investment by its mate, and will also shed light on how the pair shares incubation duties after the manipulation ends (i.e. whether the manipulation and the partner’s response influence subsequent care).

Here, we experimentally investigated the response of semipalmated sandpiper (*Calidris pusilla*) parents to the temporary absence of a partner during incubation. Semipalmated sandpipers are small shorebirds (23-32 g) that breed in the Arctic. They are considered obligate biparental incubators, i.e. the participation of both parents is required for successful incubation (Hicklin & Gratto-Trevor 2010). Incubating semipalmated sandpipers rarely feed during their incubation bout and attend their nest 95% of the time (Bulla *et al.* 2014; Bulla *et al.* 2015b). The incubation bouts last on average 11.5 hours for females and 10.7 hours for males (Bulla *et al.* 2014).

In the middle of the incubation period, we removed a parent at the end of its regular incubation bout and released it 24 hours later. In this way, the temporarily widowed bird became responsible not only for its own incubation bout (control period), but also for the following ‘incubation bout' of its partner (treated period). We investigated the change in nest attendance between control and treated period, assessed how variable this change was between individuals, and whether it was sex-dependent.

We anticipated four possible scenarios for how the temporarily widowed parent would respond (Figure 1). The Jones, Ruxton & Monaghan (2002) model predicts either no compensation (Figure 1a) or full compensation (Figure 1b), but we also consider two scenarios of partial compensation that differ in how the nest attendance changes over time (Figure 1c, d). Notably, an observation of no compensation (Figure 1a) may either reflect a decision by the bird not to change its investment in response to the absence of its partner, or a lack of knowledge about the partner’s absence (the bird may simply leave to forage at the end of its bout, as it typically does, without noticing the partner’s absence). Full compensation (Figure 1b) may arise if the incubating bird waits for the partner to return before ending its own bout, assuming that it has not yet reached its energetic limits. Partial compensation may arise when the partner continues incubation, but leaves when its energy reserves are depleted, that is, it goes from full to no compensation within its partner’s ‘bout’ (Figure 1c). However, partial compensation could also arise if the individual continues incubating, but with a lower overall nest attendance, because it regularly leaves the nest for short feeding bouts, that is, it starts behaving like a species that incubates uniparentally (Figure 1d).

**Figure 1.**
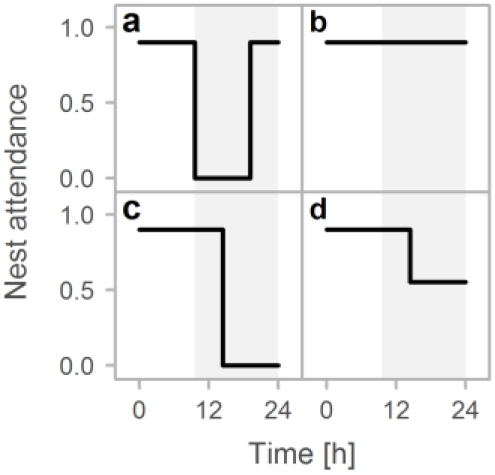
Possible compensation strategies for temporal absence of the partner. Zero represent the time when the partner was experimentally removed. The grey area represents the expected partner’s incubation bout, that is the treated period. a, No compensation – an individual leaves the nest at the end of its incubating bout and returns when its next bout is supposed to start. b, Full compensation – an individual continuous incubating for the entire supposed bout of its partner. c, Partial compensation – an individual continues incubating, but gives up and leaves at a certain point in time. d, Partial compensation – an individual continues incubating, but with lower nest attendance as it leaves the nest for short feeding bouts.

We further tested whether the following three factors might explain diversity in the compensation response. (a) Time of day: because it is less energetically demanding to incubate when it is warm (Norton 1973; Vleck 1981; Kersten & Piersma 1986; Williams 1996; Nord, Sandell & Nilsson 2010), we predict a higher level of compensation during the day compared to the colder night hours. (b) Incubation share prior to removal: individuals that have invested less in parental care in the current breeding attempt compared to others may be either more reluctant to compensate or they may have more resources left for compensation (Bowman & Bird 1987; Duckworth 1992). (c) Individual responsiveness, measured as median escape distance from the nest upon approach of a human: individuals that stay on the nest for longer when approached by a human may be less responsive and hence less likely to detect the absence of the partner or less prone to compensate (Koolhaas *et al.* 1999; Coppens, de Boer & Koolhaas 2010).

Lastly, to investigate how the experiment influenced subsequent parental care, we explored the incubation pattern of the two parents after the removed parent returned. Specifically, we investigated how nest attendance, length of incubation bouts and probability and length of exchange gaps differed before and after the experiment.

## MATERIALS AND METHODS

### Study site

We conducted the experiment in a population of semipalmated sandpipers near Barrow, Alaska (71.32° N, 156.65° W), between 1 June and 4 July 2013. The study area and species are already described in detail elsewhere (Ashkenazie & Safriel 1979a; Bulla *et al.* 2014). Barrow has continuous daylight throughout the breeding season, but environmental conditions such as ambient temperature show consistent and substantial diurnal fluctuations. Ambient temperatures are generally low, below 5 °C, but surface tundra temperatures can reach up to 28 °C (Bulla *et al.* 2014, Supplementary Figure S1)

### Recording incubation

The general procedure of monitoring incubation is described in detail elsewhere (Bulla *et al.* 2014; Bulla *et al.* 2015a). In short, the presence of parents, equipped with a plastic flag containing a glass passive tag (9.0 mm × 2.1 mm, 0.08 7g, http://www.biomark.com/), was registered every 5 s by a custom made radio frequency identification device (RFID) with a thin antenna loop around the nest cup connected to a reader. Incubation was further determined by comparing nest temperature, measured with a high resolution temperature-probe, and surface tundra temperature, measured by an MSR^®^ 145 data logger placed next to the nest. In addition, 12 nests were video recorded for some days. Fifteen out of 29 experimental nests were protected against avian predators using enclosures made of mesh wire (Supplementary Figure 1a).

### Experimental procedure

At 29 nests we temporarily removed one parent (henceforth, the ‘removed parent’) around the 11^th^ day of the 19-21 day incubation period, shortly before we expected its partner (henceforth, the ‘focal parent’) to return to incubate. We assessed the expected return of the partner by downloading the RFID data and visualizing the incubation pattern from the previous days. Alternatingly, we caught the male or the female at a nest with a spring trap triggered from a distance by a fishing line, starting with the male at the first experimental nest. The sex of individuals was known from previous years, or estimated from body measurements and later confirmed by molecular analyses using DNA extracted from a ca. 50 µl blood sample taken from the brachial vein at first capture (Bulla *et al.* 2014; Bulla *et al.* 2015a). All 29 removed individuals were at least one year old with body mass at capture between 22g and 29.7g (median 25g).

After 24 hours we released the removed parent in the vicinity of its nest. In this way, the focal parent incubated its ‘natural’ incubation bout (henceforth, the ‘control period’), which at this stage of incubation typically lasts about 10-11 hours (Bulla *et al.* 2014). The remaining 12-13 h during which the partner was removed was then considered the ‘treated period’ of the focal parent.

Nest attendance, defined as the proportion of time a bird was sitting on the nest, was derived from temperature data (Bulla 2014; Bulla *et al.* 2014), except in one nest where temperature measurements failed. In this case, attendance was derived from RFID readings because temperature-based and RFID-based attendance highly correlate: *r* = 0.79, *N* = 1584 incubation bouts from 2011 (Bulla *et al.* 2013; Bulla *et al.* 2014).

Four nests were excluded for analyses of compensation, because (a) a focal parent deserted the nest prior to treatment (one nest), (b) depredation (two nests) and (c) the wrong bird was removed (one nest). Thus, 25experimental nests with 12 females and 13 males as focal parents were used in the analyses.

### Captivity conditions

The removed parent was kept in a cardboard box (21 cm × 30 cm × 25 cm) in a shed which was sheltered from rain and wind (Supplementary Figure 1b). The bottom of the box was lined with tundra (fresh for every bird) and contained fresh water and a feeding tray (Supplementary Figure 1c).

The first eight removed birds were starved for 12 hours, and provided with ad libitum food for the remaining 12 hours. This was part of a different project and done to estimate mass loss during an incubation bout. However, after two females died on this regimen, the remaining birds were kept on ad libitum food throughout (and no further birds died).

Initially, the ad libitum food consisted of 100 mealworms (∼7.5 g) per 12 hours. The energetic content that birds can metabolize from mealworms is approximately 24.2 kJ/g (Bell 1990). Thus, ∼7.5g of worms provided ∼181 kJ, which should be more than adequate to cover the estimated daily energetic requirement of a semipalmated sandpiper during the incubation period (19-59 kJ/day; Ashkenazie & Safriel 1979b), or while resting (123 kJ/day using Norton’s (1973) equation for resting metabolic rate, assuming a 27 g bird, a median temperature of 6.2 °C and assuming that a one liter oxygen consumption is the equivalent of 20.1 kJ). However, the first two captive birds ate nearly all mealworms. Hence, we supplemented mealworms with cat food (for six birds, four of which did not eat it) or increased the amount of mealworms to 125-200 per 12 hours (for all remaining birds).

We checked and weighed each individual after capture, after 12 hours and at the end of the 24-hour captive period.

### Statistical analyses

#### Compensation for absence of the partner

We assessed whether and how the focal parent compensated for the absence of its partner by comparing nest attendance between the control and the treated period. We defined the control period as the period starting with the arrival of the focal bird on the nest (after removal of its partner) and lasting for the length of the median bout, estimated from the three before-experimental incubation bouts of the focal bird. The treated period is then the time between the end of the control period and the release of the removed partner.

To test for the difference in nest attendance between control and treated period we used linear mixed-effect models with nest attendance as the dependent variable and period (control or treated) as a categorical predictor. To account for the paired (within-individual) design of the experiment, we included bird ID as random intercept.

Nest attendance may differ depending on the length of the control or treated period (referred to as ‘period length’), but controlling for this period length did not improve the model fit; the model with period length was half as likely as the simple model (Supplementary Table 1). Hence here, and in the subsequent analyses, we made the inference from the simpler models without period length.

Next we tested whether the amount of compensation was sex-specific by comparing a model with period (control or treated) in interaction with sex, with the initial model without sex (Supplementary Table 1).

#### Explaining the diversity in compensation

To explore potential drivers of the diversity in compensation, we used linear models to test whether nest attendance during the treated period depended on (a) the time of day (defined as mid-point of the treated period, transformed to radians and represented by a sine and cosine), (b) escape distance from the nest upon approach of a human, estimated as median escape distance for an individual prior to the experiment (see Bulla *et al.* 2016), and (c) the proportion of time the focal bird was incubating before we removed its partner (estimated as median share of daily incubation, without exchange gaps, during three days prior to treatment).

We also assessed the relative importance of these three variables by comparing the three models (Supplementary Table 2). We used nest attendance during the treated period (instead of compensation), because attendance correlated strongly with compensation (*r*_Pearson_ = 0.997, *N* = 25 nests), and because the results are then directly comparable to those from the analysis of nest attendance during the control period and under natural conditions (Supplementary Table 3).

For one individual we had no escape distance estimate, so we imputed this value (following the procedure outlined in Nakagawa & Freckleton 2011) as the median of 1,000 imputations generated by the ‘amelia’ function in the ‘Amelia’ R package (Honaker, King & Blackwell 2011) with the range of likely escape distances (0 – 80 m) as a prior.

#### After-experimental effects

We explored how incubation changed after the removed parent returned to the nest (after-experimental period) by comparing – for each parent – nest attendance and the length of the last three before-experimental incubation bouts with the first three after-experimental bouts. For these bouts we also compared the presence and length of exchange gaps. To this end, we constructed mixed-effect models with nest attendance (proportion), bout length (in hours), presence of exchange gap (binomial response; 0 = no gap, 1 = gap present), and length of exchange gap (in minutes) as separate response variables, and period (before or after experiment) in interaction with day in the incubation period (day) as predictors. Day was mean-centred within each nest, so that negative values represent the before- and positive values the after-experimental period. To control for non-independence of data we entered bird ID as a random intercept and day as a random slope. We assessed the importance of the interaction and of the type of parent (focal versus removed) by comparing models with and without the interaction, and with and without parent type (Supplementary Tables 4 and 5).

In addition, we explored whether the after-experimental nest attendance and bout length of the removed parent were related to its mass loss while in captivity, and whether the after-experimental nest attendance and bout length of the focal parent were related to its level of compensation (proportion) during the treated period. We also investigated whether these relationships were sex specific. Bird ID was entered as a random intercept, and mass loss or compensation as random slopes (Supplementary Table 6 and 7).

#### General procedures

R version 3.3.0 (R-Core-Team 2016) was used for all statistical analyses and the ‘lme4’ R package (Bates *et al.* 2015) for fitting the mixed-effect models. The models were fitted with maximum likelihood. We used the ‘sim’ function from the ‘arm’ R package and non-informative prior-distribution (Gelman & Hill 2007; Gelman & Su 2015) to create a sample of 2,000 simulated values for each model parameter (i.e. posterior distribution). We report effect sizes and model predictions by the medians, and the uncertainty of the estimates and prediction by the Bayesian 95% credible intervals represented by 2.5 and 97.5 percentiles (95%CI) from the posterior distribution of the 2,000 simulated or predicted values. We estimated the variance components with the ‘lmer’ or ‘glmer’ function from the ‘lme4’ R package (Bates et al. 2015).

By necessity, the dependent variables varied more for the treated or the after-experimental period than for the control or before-experimental period. We controlled for this heteroscedasticity by scaling the dependent variable within period. However, because these models generated similar results as the simpler models and because the simpler models are on the original scale and hence easier to interpret, we report only the outcomes of the simpler models.

In all model comparisons we assessed the model fit by Akaike’s Information Criterion corrected for sample size (AICc; Anderson 2008) generated by the ‘AICc’ function from the ‘AICcmodavg’ R package (Mazerolle 2016).

### Ethical statement

All field procedures were performed in accordance with the relevant guidelines and regulations, and approved by the U.S. Department of the Interior, U.S. Fish and Wildlife Service, and State of Alaska Department of Fish and Game (permit no. 23520, MB210494-2, 13-122).

## RESULTS

### Compensation for absence of parental care

Typically, parents partially compensated for the absence of their partner’s care (Figure 2a). The nest attendance (proportion of total time on the nest) in the treated period was on average 0.38 (95%CI: 0.27-0.49) lower than in the control period (Figure 2b; Supplementary Table 1). This translates to a 59% (49-70%) compensation for the absence of the partner. The level of compensation was similar for females and males (Figure 2b, Supplementary Table 1; the model containing sex in interaction with treatment was less likely than the model without this interaction).

The compensation response of individual parents ranged from no to full compensation (range: 2-101%, median = 57%; Fig. 2a). Birds achieved similar levels of partial compensation using various ‘strategies’ (Figure3). Some individuals gradually decreased their nest attendance over the experimental period; some compensated fully for part of the experimental period, but then either reduced their nest attendance, left the nest completely unattended, or left the nest unattended but came back later. Remarkably, the individuals with nearly no compensation during the treated period simply returned to the nest at the expected time for their next incubation bout, that is, they continued their pre-experimental incubation routine (Figure 3, top row). In contrast, the parents that fully compensated left the nest unattended after continuously incubating for more than 24 hours (Figure 3, panels in two bottom rows).

**Figure 2.**
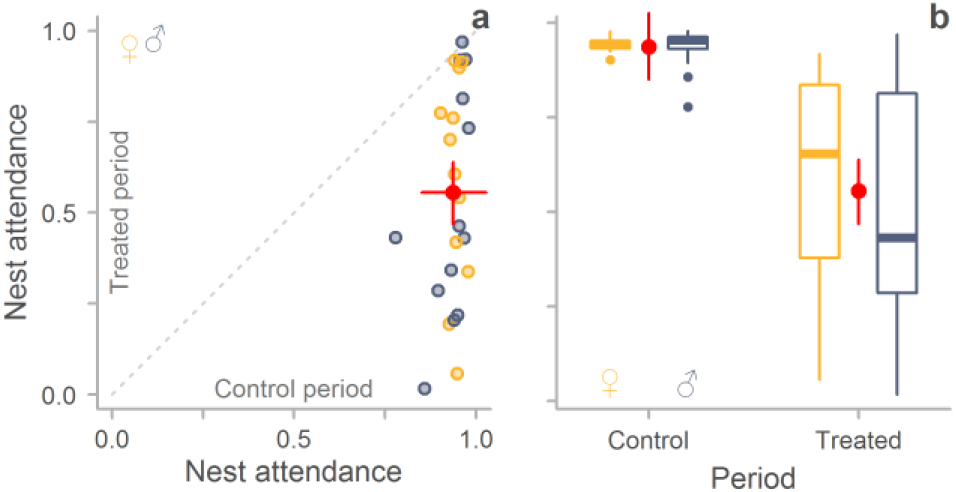
Compensation for the temporary absence of the partner. a-b, The control period reflects the regular incubation bout of the focal parent, while the treated period is the period during which the focal parent’s partner should have incubated, but was held in captivity. Yellow indicates female, grey-blue male. Red dots with bars indicate model predictions with 95%CI (Supplementary Table 1). a, Compensation by each individual (*N* = 25). The grey dashed line indicates full compensation. Each dot represents the nest attendance of one individual; points below the line represent various degrees of partial compensation or no compensation (zero). b, Nest attendance in the control and treated period for females (*N* = 12) and males (*N* = 13) separately. Box plots depict median (horizontal line inside the box), the 25^th^ and 75^th^ percentiles (box), the 25^th^ and 75^th^ percentiles ±1.5 times the interquartile range or the minimum/maximum value, whichever is smaller (bars), and the outliers (dots).

**Figure 3.**
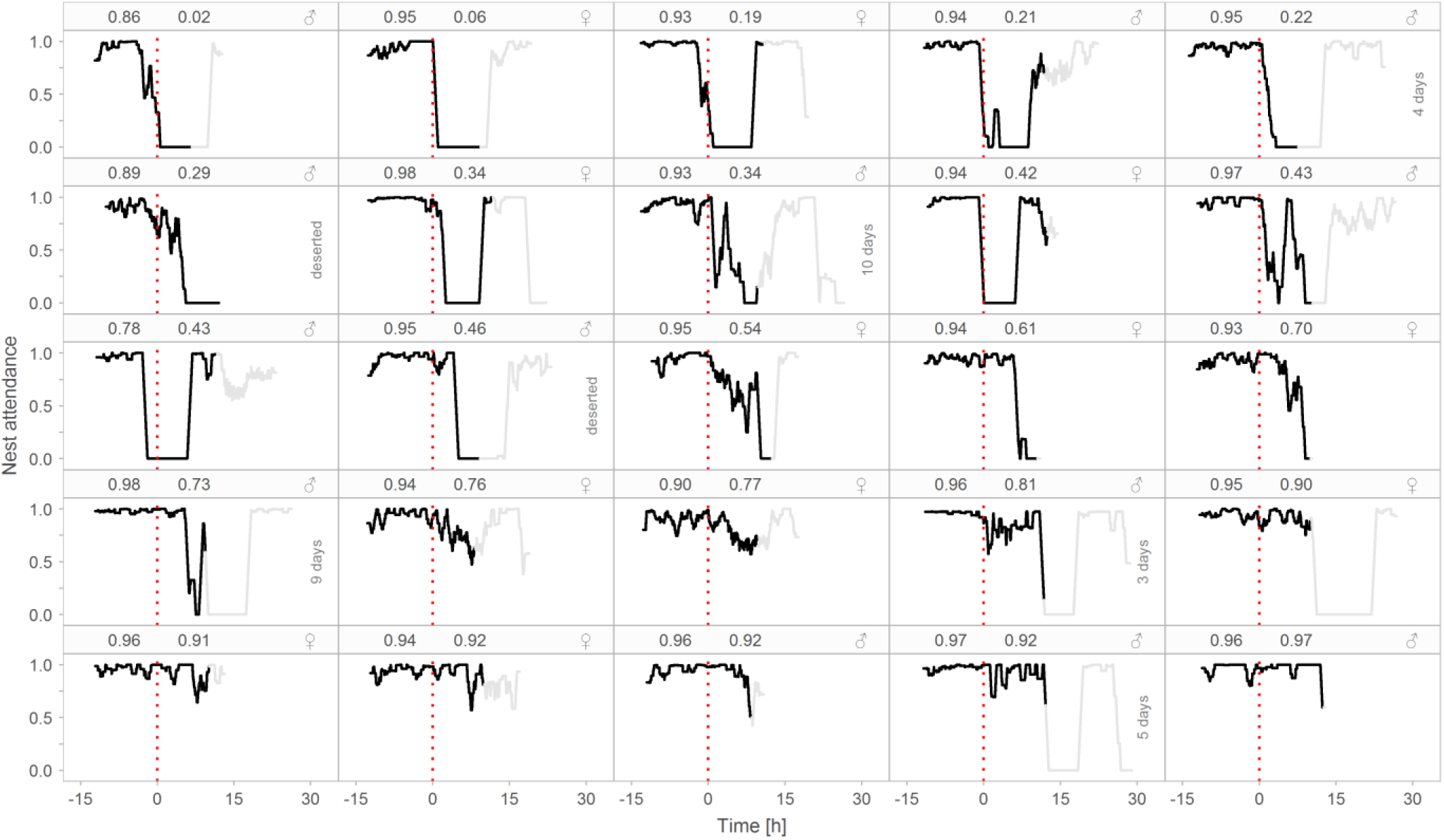
Diverse compensation responses by individual parents in the absence of their partner. Each panel represents one of 25 focal individuals. Panels are ordered according to nest attendance within the treated period such that the individual with the lowest nest attendance is in the top-left panel. Black lines show hourly nest attendance (proportion of time the parent is on the nest, depicted as a running mean) during the experimental period (i.e. from the return of the focal parent until the release of the removed parent). The red dotted lines (time zero) indicate the end of the control period (i.e. the regular incubation bout of the focal parent; negative values) and the start of the treated period (compensation period, positive values). Grey lines indicate the hourly nest attendance of the focal bird from the moment the removed parent was released until it returned to the nest. In seven nests the removed parent never returned, so we show a maximum of 30 hours after the start of the treated period and note whether the incubating parent deserted within this period, or for how many days the individual continued incubating uniparentally.

### Explaining the diversity in compensation

Whereas during undisturbed situations (the before-experimental period or non-experimental nests) nest attendance slightly decreased during the warmer part of the day (grey-blue and green in Figure 4a), the parents that compensated for the absence of their partner during the warmer part of the day tended to have higher nest attendance (yellow in Figure 4a). Compensation seemed unrelated to the focal parent’s share of incubation (proportion) during the before-treatment period (Figure 4b), but there was a tendency for parents with long escape distance (i.e. parents sensitive to disturbance) to compensate more (Figure 4c). The escape distance model had the greatest support of the three models (see Supplementary Table 2).

**Figure 4.**
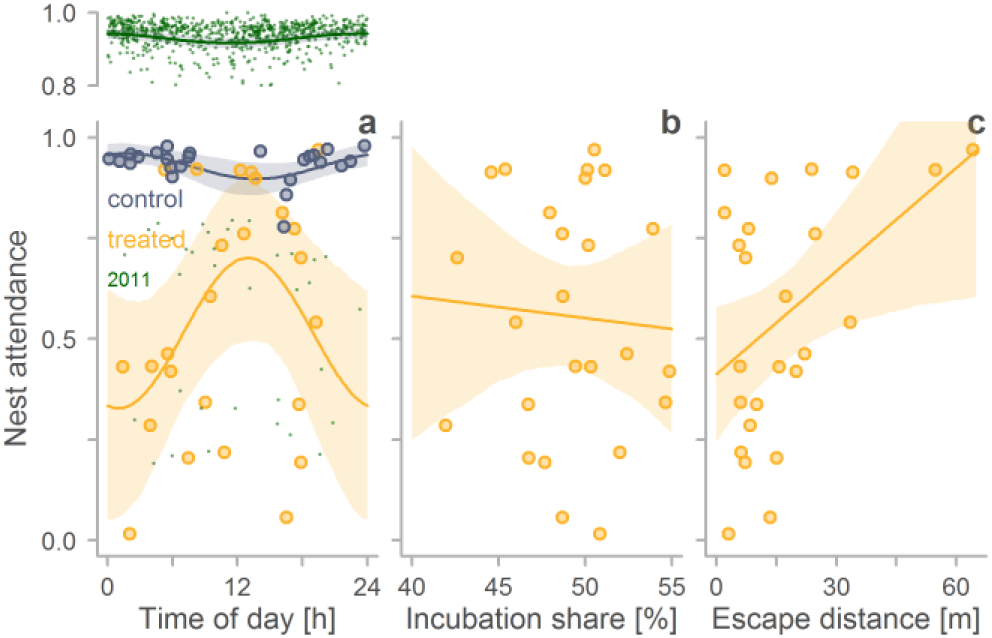
Correlates of compensation for the absence of the partner. a-c, Relationships between nest attendance (proportion of the total time the bird is on the nest) during the time the removed partner would have incubated (treated period) and the mid-time of this treated period (a), the focal parent’s share of incubation during the before-experimental period (b) and its escape distance from the nest prior to the experiment (c). Yellow dots represent individual observations (*N* = 25); yellow lines with yellow-shaded areas indicate model predictions with 95%CI for the treated period (Supplementary Table 2). a, To emphasize how the relationship of nest attendance with time of day differs between the treated period and the natural, undisturbed situation, we added the observations and predictions from the control period (i.e. the regular incubation bout of the focal parent; grey-blue) and from non-experimental nests from the 2011 breeding season (green, Supplementary Table 3; the 2011 data come from Bulla *et al.* 2013; Bulla *et al.* 2014).

### After-experimental effects

In five out of 25 experimental nests the removed parent (all females) never returned to the nest after release. In an additional two nests the removed females never returned, but these nests were excluded from the main analyses, because one was partially depredated during the incubation bout prior to removal, while in the other the focal bird (male) had already deserted the nest before the treated period started.

The five widowed males and the additional two experimentally widowed males (at different nest than described above; see Methods) continued incubating for another 0-10 days (median = 4 days). They then deserted the nest (*N* = 5), possibly hatched one egg and deserted the remaining three eggs (*N* = 1), or the nest was depredated (*N* = 1).

**Figure 5.**
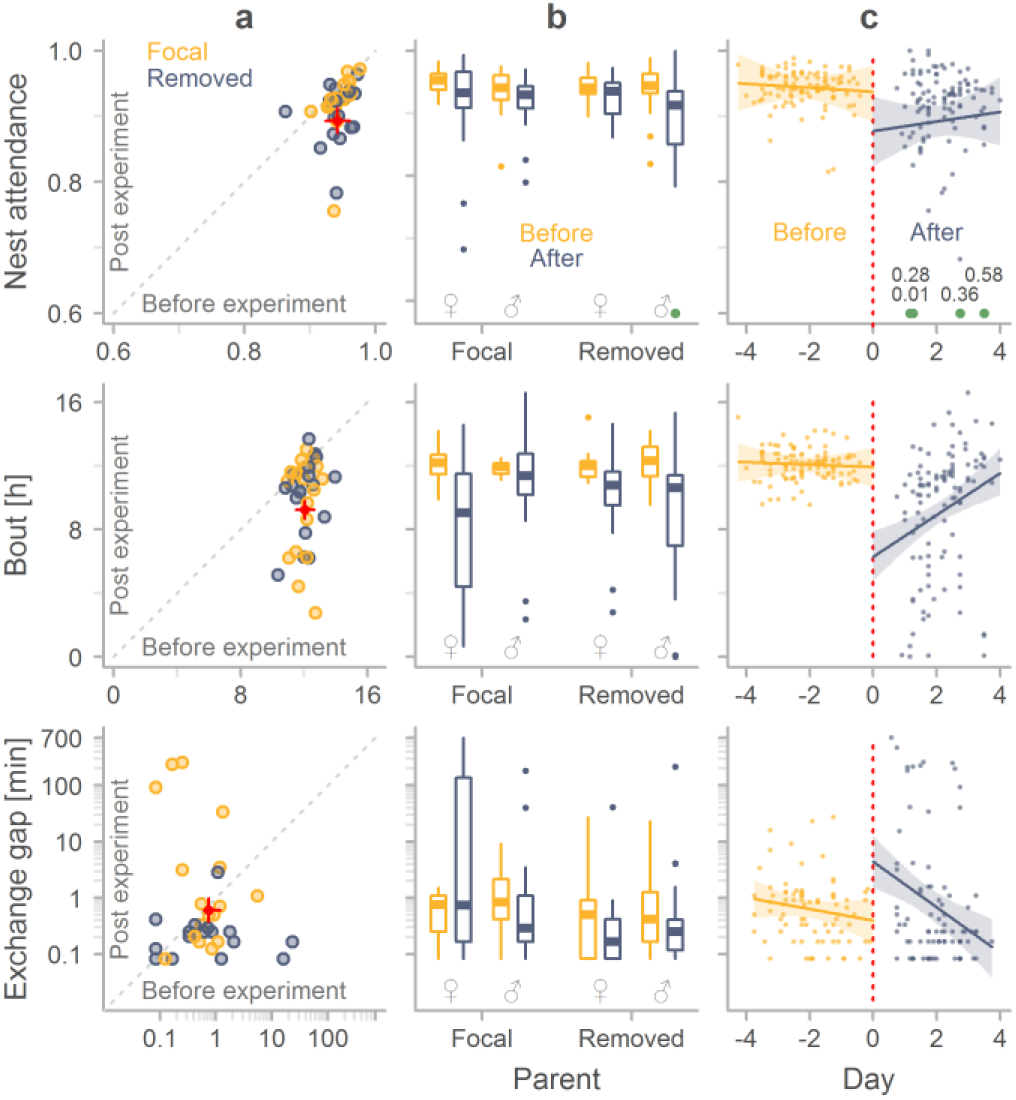
Differences in quality of incubation before and after the experimental period. a, Median nest attendance, median bout length and median non-zero exchange gap duration for each individual in the period before and after the experiment. Dots represent medians for focal parents (yellow) and for removed parents (grey-grey-blue; *N*_nest attendance_ and *N*_bout_ = 36 individuals, *N*_gap_ = 33 individuals). Red dots with bars indicate model predictions with 95%CI (Supplementary Table 4 and 5, ‘simple model’). b, Comparison of nest attendance, bout length and exchange-gap length between the period before (yellow) and after (grey-grey-blue) the experiment for the focal and the removed parent and for each sex separately (*N*_nest attendance_ and *N*_bout_ = 214, *N*_gap_ = 164; for description of boxplots see legend Figure 2). For nest attendance, the green dot represents four outliers (described in c). c, Temporal changes in nest attendance, bout or gap length in the period before (yellow, negative values) and after (grey-grey-blue, positive values) the experiment (focal and removed individuals combined). Dots represent individual observations and lines with shaded areas indicate model predictions with 95%CI (Supplementary Table 4 and 5, ‘day model’). See b for sample sizes. The red dotted line indicates the day when one of the parents was removed. In the nest attendance graph, four values are outside the range of the y-axis; these are indicated in green with their actual nest attendance value. In case of nest attendance the model including day was slightly less likely than the simple model. In case of bout length and exchange gap duration, the model including day was much more likely than the simple model. For exchange-gap duration, the model that also contained parent type (focal or removed) was even more likely (Supplementary Tables 4 and 5).

In the 18 nests where the removed parent returned to incubate, parents differed markedly in how long it took them to return: median (range) = 7.36 hours (0.26-16.85 hours). In these 18 nests, the overall quality of incubation during the after-experimental period was lower than during the before-experimental period (Figure 5; Supplementary Tables 4 and 5): nest attendance was lower, incubation bouts were shorter and exchange gaps, although they did not occur more frequently, were longer (Figure 5a). These effects were similar between focal and removed parent (Figure 5b, Supplementary Tables 4 and 5), as well as between males and females (Figure 5b). However, parents seemed to recover from the effect of the treatment, because nest attendance tended to increase, bouts became longer, and gaps shortened with days after the experimental period (Figure 5c).

During the after-experimental period, nest attendance tended to be lower and incubation bouts shorter in males (but not in females) that lost more mass while in captivity (Figure 6a, Supplementary Table 6). Although the level of compensation seemed unrelated to nest attendance in after-experimental bouts, females (but not males) that compensated more tended to have shorter bouts (Figure 6b, Supplementary Table 7).

**Figure 6.**
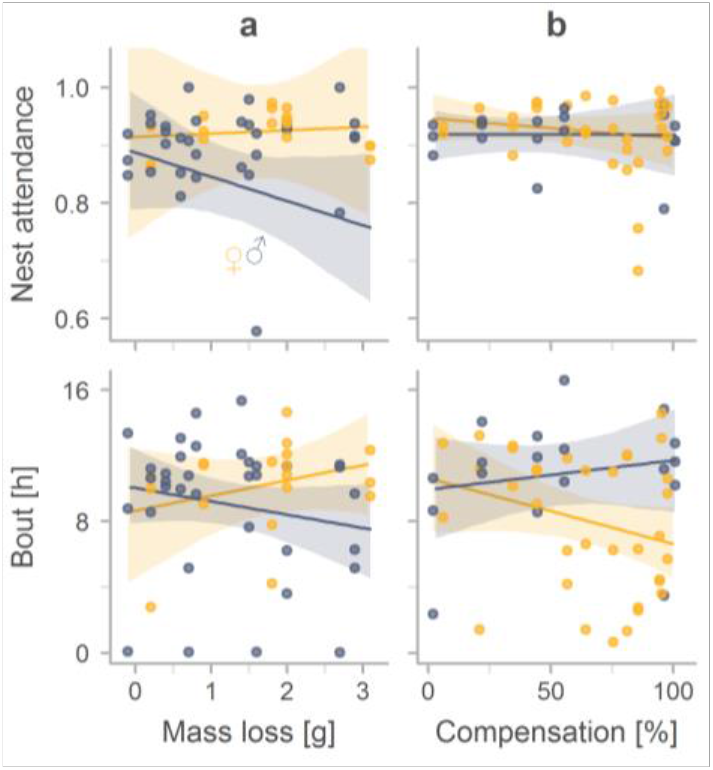
Predictors of nest attendance and bout length during the after-experimental period. a, Relationship between mass loss of the removed parent while in captivity and its nest attendance and bout length during the post-experimental period. b, Relationship between the amount of compensation by the focal parent during the time the removed partner would have incubated (treated period) and the focal parent’s nest attendance and bout length during the after-experimental period. a-b, Yellow indicates females, grey-grey-blue males; dots represent individual observations and lines with shaded areas indicate model predictions with 95%CI. In all four cases the model with sex fitted the data better (Supplementary Tables 6 and 7).

## DISCUSSION

### Diverse compensation

Our results indicate that semipalmated sandpipers on average partially compensate for the temporal absence of care from their partner. This contradicts the prediction of the incubation model (Jones, Ruxton & Monaghan 2002), but seems in line with the general predictions of established parental care models (Houston & Davies 1985; McNamara, Gasson & Houston 1999; McNamara *et al.* 2003). However, parents varied greatly in how they responded (Figure 3): some parents did not compensate at all, some showed various degrees of partial compensation, and some compensated fully. We discuss two possible explanations for this diversity.

In semipalmated sandpipers, and in many other biparentally incubating shorebirds, the contribution of both parents is thought to be essential for successful incubation (Poole 2005). Thus, according to theory, parents faced with a temporal decrease in incubation from their partner should either fully compensate or desert the nest (Jones, Ruxton & Monaghan 2002). However, a reduction in incubation effort may not always lead to complete loss of the breeding attempt. For example, during warmer periods, near the end of the incubation period, or when a parent has larger energy reserves, a single parent might successfully incubate a clutch (Bulla *et al.* 2017). This means that there might be some room for one parent to exploit the investment of the other parent. Indeed, one permanently widowed parent incubated uniparentally for 10 days (see Actograms in Supporting Information (Bulla 2017a) from Bulla *et al.* 2017) and some non-experimental nests hatched after 14 days of uniparental incubation (Bulla *et al.* 2017). Thus the varying circumstances among nests could translate into various compensation levels.

Alternatively, semipalmated sandpiper parents may always attempt to compensate fully, but sometimes fail to do so, either (a) because their energy reserves get depleted (Figure 3), or (b) because they are less responsive to the absence of their partner’s presence at the nest (Figure 4c).

(a) Parents of biparental species with continuous incubation (i.e. with close to 100% nest attendance) starve while incubating (Chaurand & Weimerskirch 1994; Weimerskirch 1995; Dearborn 2001; Bulla *et al.* 2014; Bulla *et al.* 2015b). Thus, parents are obviously unable to incubate continuously for many days. This implies that full compensation is only possible as long as the energetic reserves last. We find some support for this explanation. As we demonstrate, even the parents that compensated fully left the nest unattended after some time (Figure 3), that is, they did not (could not) continue incubating for three ‘typical’ incubation bouts. Thus, the fully compensating parents differed from the not- or partially-compensating parents by ‘deserting’ the nest considerably later. Further evidence comes from the observation that parents that were treated during the warmer part of a day (when incubation is presumably less energetically demanding) tended to compensate more (i.e. had higher nest attendance) than parents treated during the colder part of a day. This contrasts with the typical nest attendance patterns when both parents are present (Figure 4), as well as with nest attendance patterns of uniparentally incubating species (Cartar & Montgomerie 1985; Løfaldli 1985; Reneerkens *et al.*2011; Bulla *et al.* 2017). In both these cases, nest attendance drops during the warmer part of a day, probably because the eggs cool down slower and because food availability and hence foraging efficiency is higher. Thus, our results are consistent with the idea that the compensating parents might have tried but could not compensate fully when it was cold.

(b) An alternative idea is that the level of compensation depends solely on the perceived absence of the partner’s nest attendance. Unlike chick feeding, where parents can feed simultaneously, incubation is a mutually exclusive behaviour, because only a single parent can incubate at a time. However, the off-nest parent is often far away from the nest, clearly out of hearing range of its partner (Bulla *et al.* 2015b). Observations show that the incubating parent sometimes leaves the nest before its partner returns to incubate (Ashkenazie & Safriel 1979b; Bulla *et al.* 2014; Bulla *et al.* 2015b), and may ‘assume’ that its mate will return and continue incubation. Variation in compensation may then be related to the variation in how often or how soon the partner checks its nest, or to variation in how long a parent waits for its partner to return, that is, the responsiveness to the partner’s absence (Figure 4c). Indeed, some permanently widowed parents continued their typical incubation schedule, leaving the nest unattended during their partner’s supposed bout, for several days before changing to a uniparental incubation pattern, suggesting that it took some time before they realized that their partner had deserted or at least before they responded to it (see Actograms in Supporting Information (Bulla 2017a) from Bulla *et al.* 2017). This also fits with our observation that more responsive individuals – i.e. those with a longer escape distance – compensated more during the experimental period than those that were less responsive, i.e. had a shorter escape distance (Figure 4c).

Energetic constraints and responsiveness may well act together. Thus, those parents that are responsive to the absence of their partner, and have the resources to wait for their partner’s delayed return, may do so, whereas parents that are less responsive or do not have the resources for full compensation, may compensate partially or not at all. Such an explanation is in line with predictions of parental care models: parents should vary in their compensation response based on the likelihood of brood failure in the absence of care, the parent’s current condition and their knowledge about (or – as we suggest – their responsiveness to) their partner’s condition or the need of the brood (Jones, Ruxton & Monaghan 2002; Johnstone & Hinde 2006). In this case, the need of the brood can be translated to the risk of temperature-related embryo death (or developmental problems affecting future fitness) or the risk of clutch predation.

### After-experimental effects

After release from captivity, 5 out of 25 parents never returned to incubate. In all cases, the non-returning parent was the female of the pair, which is similar to what has been shown in northern flickers, *Colaptes auratus* (Wiebe 2010). Females might be more sensitive to stress, because they already laid the eggs (typically, a four-egg clutch is laid in five days and has a similar total mass as an average female’s body mass; Hicklin & Gratto-Trevor 2010). In semipalmated sandpipers, females also tend to desert the brood before or after hatching (Hicklin & Gratto-Trevor 2010; Bulla *et al.* 2017). However, we found no marked differences between females and males in the level of compensation during the partner’s absence (Figure 2), or in postexperimental quality of incubation (Figure 5).

After we released the removed parent the quality of incubation was lower than before the experiment, but it improved quickly with time (Figure 5); already three days after the experiment, parents seemed to incubate as usual. The after-experimental effects were generally similar for the focal and the removed parent (Figure 5b, Supplementary Tables 4 and 5), suggesting that the stress caused by the absence of the partner (including the compensation) might have been similar to the stress of captivity. An alternative explanation for the lower quality in after experimental period is that parents needed to ‘renegotiate’ how much they invest, or realign their incubation schedules.

Mass loss of the removed parent during captivity and the amount of compensation of the focal parent during the experiment were poor predictors of the after-treatment incubation behaviour (Supplementary Table 6 and 7, ‘simple models’). This suggests that the after-experimental effects are not related to energetic constraints, confirming earlier work (Bulla *et al.* 2015a; Bulla *et al.* 2016). However, we found a tendency for sex-specific effects (Figure 5), which deserve future verification.

### Conclusions and suggestions for further work

Our finding that biparentally incubating shorebirds on average partially compensate for the temporal absence of their partner corroborates the predictions of established models (Houston & Davies 1985; McNamara, Gasson & Houston 1999; McNamara *et al.* 2003) and results of a meta-analysis (Harrison *et al.* 2009). However, the individual responses were highly diverse, from no to full compensation. Whether this variation represents noise around the mean or biologically relevant diversity awaits future investigations. Because full or nearly full compensation was the most common response, we speculate that all individuals attempt full compensation, but that some fail because their energy stores get depleted, or because they are less responsive to the absence of their partner. We provide some correlational evidence for the latter. It is also possible that the responses depend on environmental factors such as temperature and food availability. Thus, one could speculate that with increasing temperatures in the Arctic due to climate change, full compensation (and hence uniparental incubation) might become more common.

We suggest an experimental approach to test whether energy reserves drive the level of compensation. If compensation depends on energetic constraints, then supplemental feeding or heating the eggs of the focal parent (see Bulla *et al.* 2015a) should lead to full compensation in all individuals, or at least to reduced individual variation in the level of compensation. On the other hand, if parental responsiveness drives the level of compensation, and if this is an individual-specific trait, then the level of compensation should be repeatable

The diversity of compensation responses during incubation, and the role of energetic constraints and individual responsiveness (‘personality’) of the parents deserve further empirical and theoretical work.

## Author’s contributions

M.B. and B.K. conceived the study; M.B. with help of A.R. collected the data; M.B. and A.R. managed the database; M.B. coordinated the study, analysed the data with help from M.V., and wrote the manuscript with help from B.K. All authors contributed to the final paper.

## Acknowledgements

We thank F. Heim, L. Verlinden, and M. Schneider for help in the field, M. Hanson and E. Burnet from UMIAQ and R. Lanctot for assistance with logistics, A. Girg for the genetic sexing, F. Korner-Nievergelt, and Y. Araya for advice on the statistical analyses, and D. Starr-Glass, B. Bulla, P. B. D'Amelio and E. Schlicht for constructive suggestions on the manuscript. M.B. thanks Bare and Maje for patience and support. This work was funded by the Max Planck Society (to B.K.) and EU Marie Curie individual fellowship (project SocialJetLag - 4231.1; to M.B.). M.B. did this work as a PhD student in the International Max Planck Research School for Organismal Biology. The authors declare no conflicts of interest.

## Data and code accessibility

All statistical analyses are replicable with the open access data and r-code that will be available from https://osf.io/mx82q (Bulla 2017b), which will also contain all supporting information.

## SUPPORTING INFORMATION

(will be online on-line only and available from Open Science Framework https://osf.io/mx82q)

**Supplementary Figure 1.**
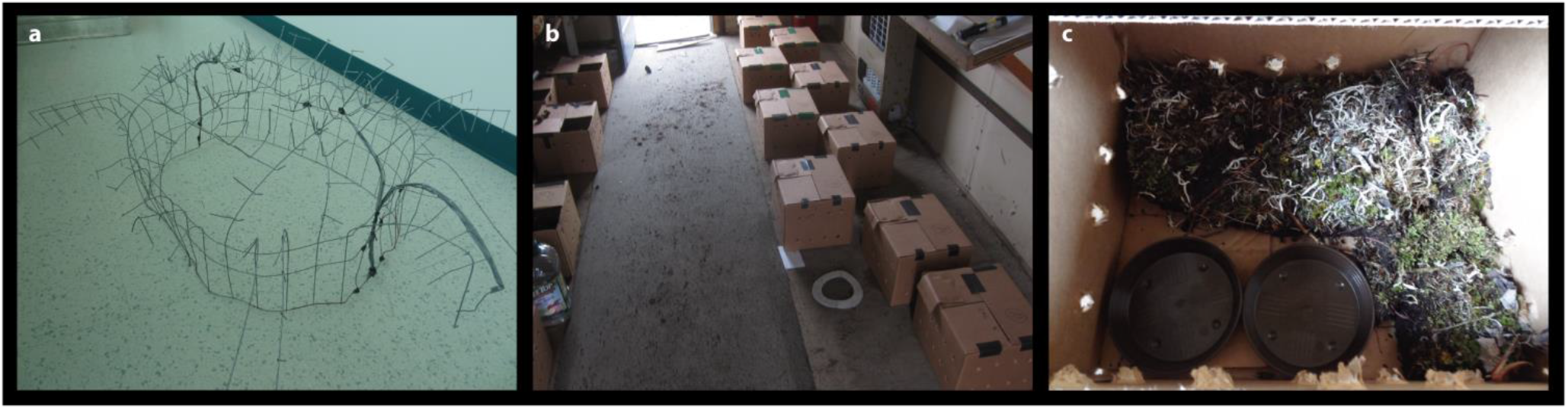
Enclosure for nest-protection, boxes and shed for removed parents. **a,** Enclosure that protected the nest against avian predators (approximate size 0.8 × 0.7 × 0.5 m) was a modification of an earlier design (Bulla *et al.* 2015a) and was made by cutting and bending a chicken wire (mesh size 5 × 5 cm and 5 × 10 cm where the cage touched the ground; wire ø 1.9 mm). The sharp parts sticking out into the air prohibit avian predators to land on the enclosure. The enclosure allowed incubating parents to fly off the nest, which was not the case for the earlier design (Bulla *et al.* 2015a) – where some birds had difficulties escaping upon approach of avian predator (we recorded one case where a parent was caught by the avian predator). **b,** Removed parents were brought to the shed where they were kept in cardboard boxes (21 cm × 30 cm × 25cm; with holes on the sides). **c,** Each box was lined with tundra and contained water and feeding tray.

**Supplementary Table 1.** Effects of treatment on nest attendance.

| Model | AIC | $\Delta AICc^a$ | $w_i^b$ | ER<sup>d</sup> | Effect type | Effect | Estimate | 95% CI | |
| --- | --- | --- | --- | --- | --- | --- | --- | --- | --- |
|  |  |  |  |  |  |  |  | Lower | Upper |
| Minimal | -5.64 | 0 | 0.624 | 1 | Fixed | Intercept (control) | 0.939 | 0.854 | 1.024 |
|  |  |  |  |  |  | Period (treated) | -0.384 | -0.494 | -0.266 |
|  |  |  |  |  | Random (variance) | Bird ID (intercept) | 9% |  |  |
|  |  |  |  |  |  | Residual | 91% |  |  |
| With period Length | -4.19 | 1.45 | 0.302 | 2.06 | Fixed | Intercept (control) | 0.915 | 0.823 | 1.011 |
|  |  |  |  |  |  | Period (treated) | -0.338 | -0.484 | -0.206 |
|  |  |  |  |  |  | Period length | 0.035 | -0.034 | 0.106 |
|  |  |  |  |  | Random (variance) | Bird ID (intercept) | 9% |  |  |
|  |  |  |  |  |  | Residual | 91% |  |  |
| With sex | -1.36 | 4.28 | 0.074 | 8.48 | Fixed | Intercept (control, ♀) | 0.942 | 0.811 | 1.063 |
|  |  |  |  |  |  | Period (treated) | -0.345 | -0.517 | -0.179 |
|  |  |  |  |  |  | Sex (♂) | -0.012 | -0.183 | 0.156 |
| | | | | | | Period × Sex | -0.066 | -0.304 | 0.165 |
|  |  |  |  |  | Random (variance) | Bird ID (intercept) | 9% |  |  |
|  |  |  |  |  |  | Residual | 91% |  |  |

**Supplementary Table 2.** Predictors of nest attendance during the treated period.

| Model | AIC | $\Delta AICc^a$ | $w_i^b$ | ER<sup>d</sup> | Effect | Estimate | 95% CI | |
| --- | --- | --- | --- | --- | --- | --- | --- | --- |
|  |  |  |  |  |  |  | Lower | Upper |
| Time | 15.71 | 4.74 | 0.08 | 11 | Intercept | 0.518 | 0.391 | 0.645 |
|  |  |  |  |  | sin (time) | -0.051 | -0.212 | 0.111 |
|  |  |  |  |  | cos (time) | -0.169 | -0.381 | 0.028 |
| Proportion | 16.74 | 5.78 | 0.05 | 18 | Intercept | 0.832 | -1.145 | 2.725 |
|  |  |  |  |  | Proportion of incubation | -0.551 | -4.411 | 3.484 |
| Escape | 10.97 | 0 | 0.87 | 1 | Intercept | 0.405 | 0.245 | 0.569 |
|  |  |  |  |  | Escape distance (m) | 0.009 | 0.001 | 0.016 |

Shown are the posterior estimates (medians) of the effect sizes with the 95% credible intervals (CI) from a posterior distribution of 2,000 simulated values generated by the 'sim' function in R (Gelman & Su 2015). $N = 25$ individuals.
<sup>a</sup>The difference in AICc between the first-ranked model and the given model.
<sup>b</sup>Akaike weight – the weight of evidence that a given model is the best approximating model (i.e., probability of the model).
<sup>d</sup>Evidence ratio – model weight of the first-ranked model relative to that of the given model (i.e., how many times is the first-ranked model more likely than the given model).

**Supplementary Table 3.** Relationship between nest attendance and time-of-day during the control period and under natural conditions.

| Model | Effect type | Effect | Estimate | 95% CI |  |
| --- | --- | --- | --- | --- | --- |
|  |  |  |  | Lower | Upper |
| Control period | Fixed | Intercept | 0.929 | 0.911 | 0.947 |
|  |  | sin (time) | 0.015 | -0.005 | 0.035 |
|  |  | cos (time) | 0.028 | -0.003 | 0.058 |
| Natural conditions<br>2011 data | Fixed | Intercept | 0.929 | 0.918 | 0.939 |
|  |  | sin (time) | 0.015 | -0.011 | 0.007 |
|  |  | cos (time) | 0.028 | 0.003 | 0.021 |
|  | Random<br>(variance) | Bird ID (intercept) | 22% |  |  |
|  |  | sin (time) | 1% |  |  |
|  |  | cos (time) | 1% |  |  |
|  |  | Residual | 76% |  |  |

Shown are the posterior estimates (medians) of the effect sizes with the 95% credible intervals (CI) from a posterior distribution of 2,000 simulated values generated by the 'sim' function in R (Gelman & Su 2015). Variance components were estimated by the 'lmer' function in R (Bates *et al.* 2015).
$N_{\text{control}} = 25$ control periods (one per focal experimental parent); control period started with the arrival of the focal bird on the nest (after removal of its partner) and lasted for the length of median incubation bout, estimated from the three pre-experimental incubation bouts of the focal bird.
$N_{\text{2011 data}} = 832$ incubation bouts from 47 nests (the data come from Bulla *et al.* 2013; Bulla *et al.* 2014)

**Supplementary Table 4.**
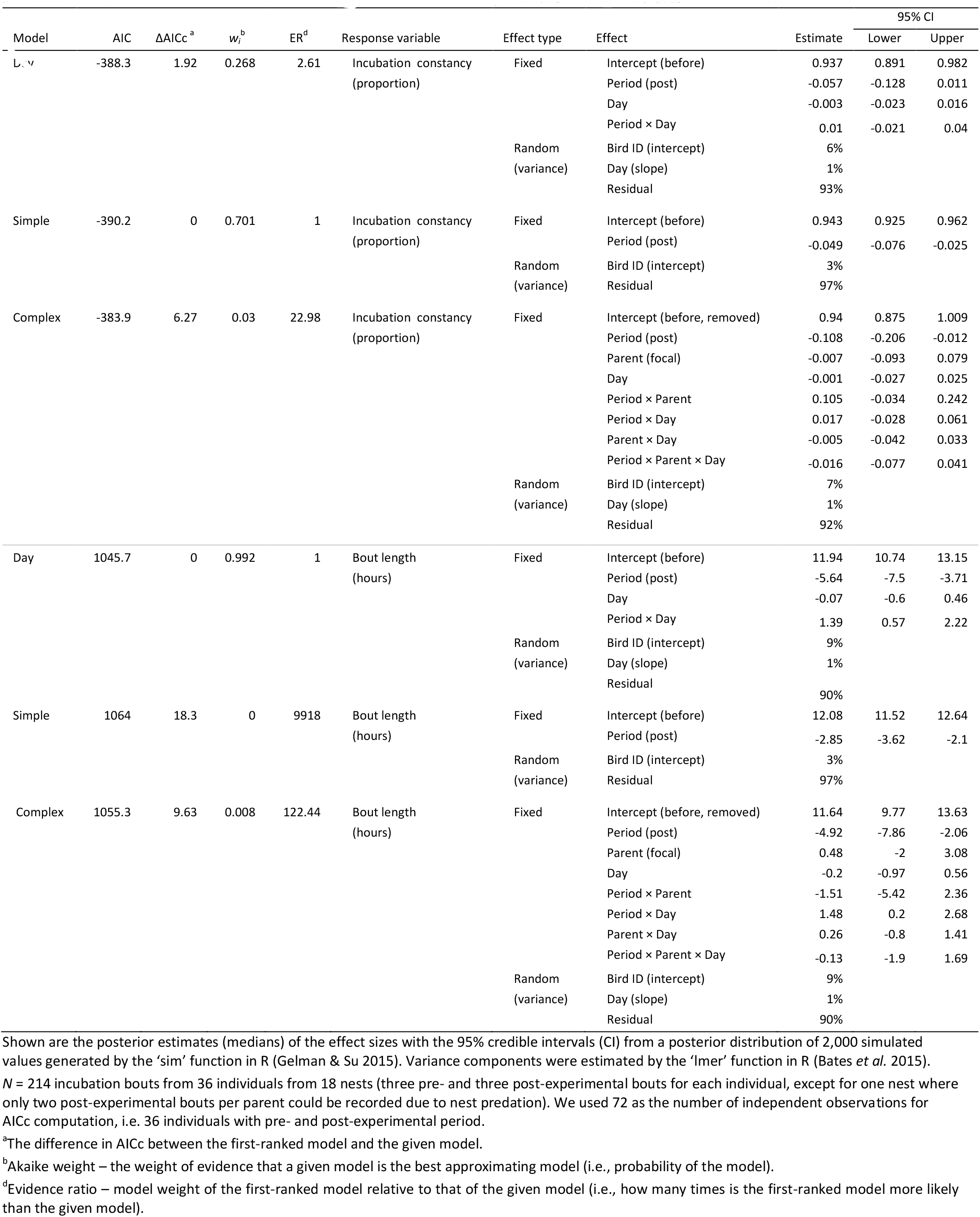
Difference in nest attendance and bout length in the before- and after-experimental period.

**Supplementary Table 5.**
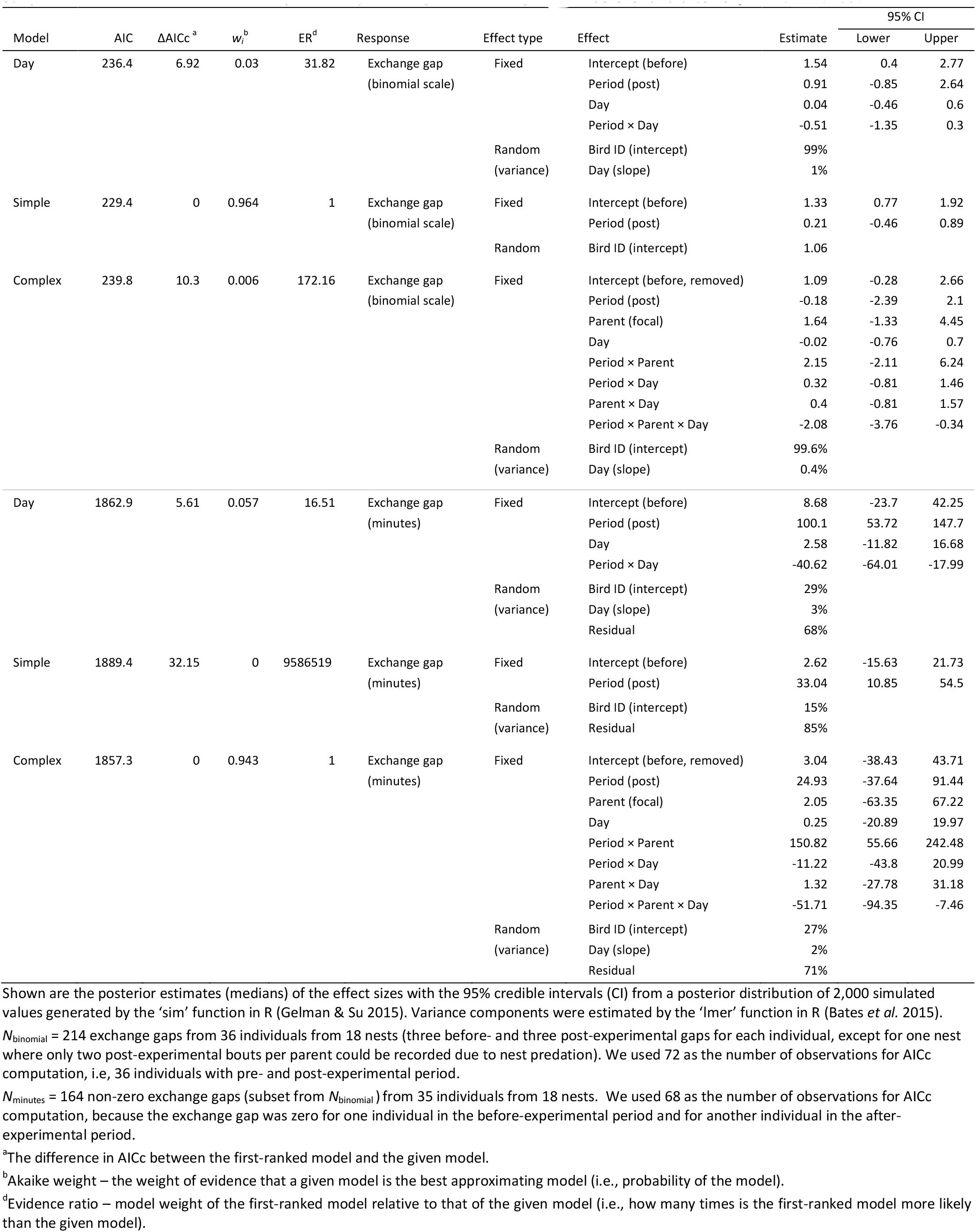
Difference in probability and length of exchange gap in before- and after-experiment period.

**Supplementary Table 6.**
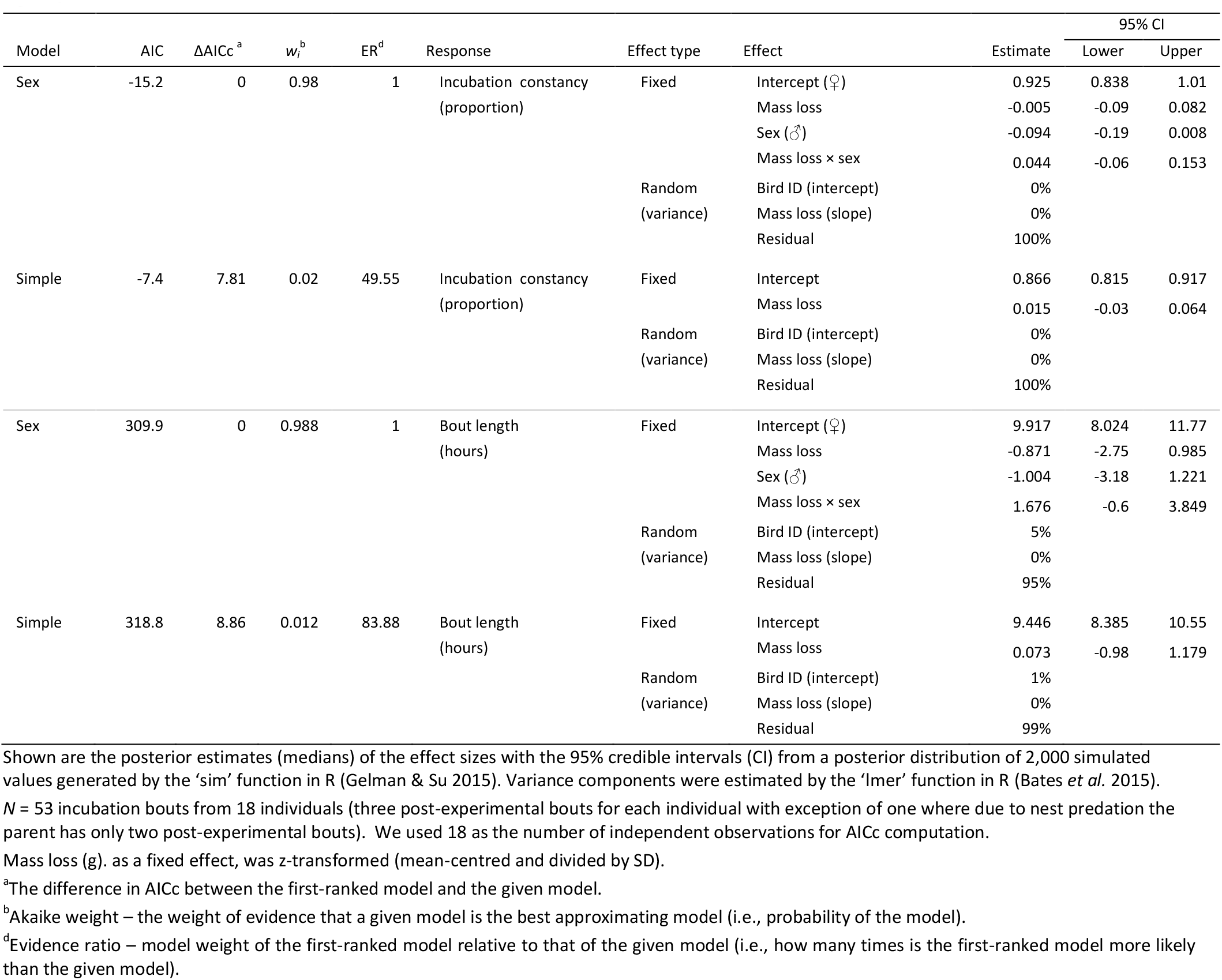
Effect of mass loss of the removed parent on its nest attendance and bout length in the after-experimental period.

**Supplementary Table 7.**
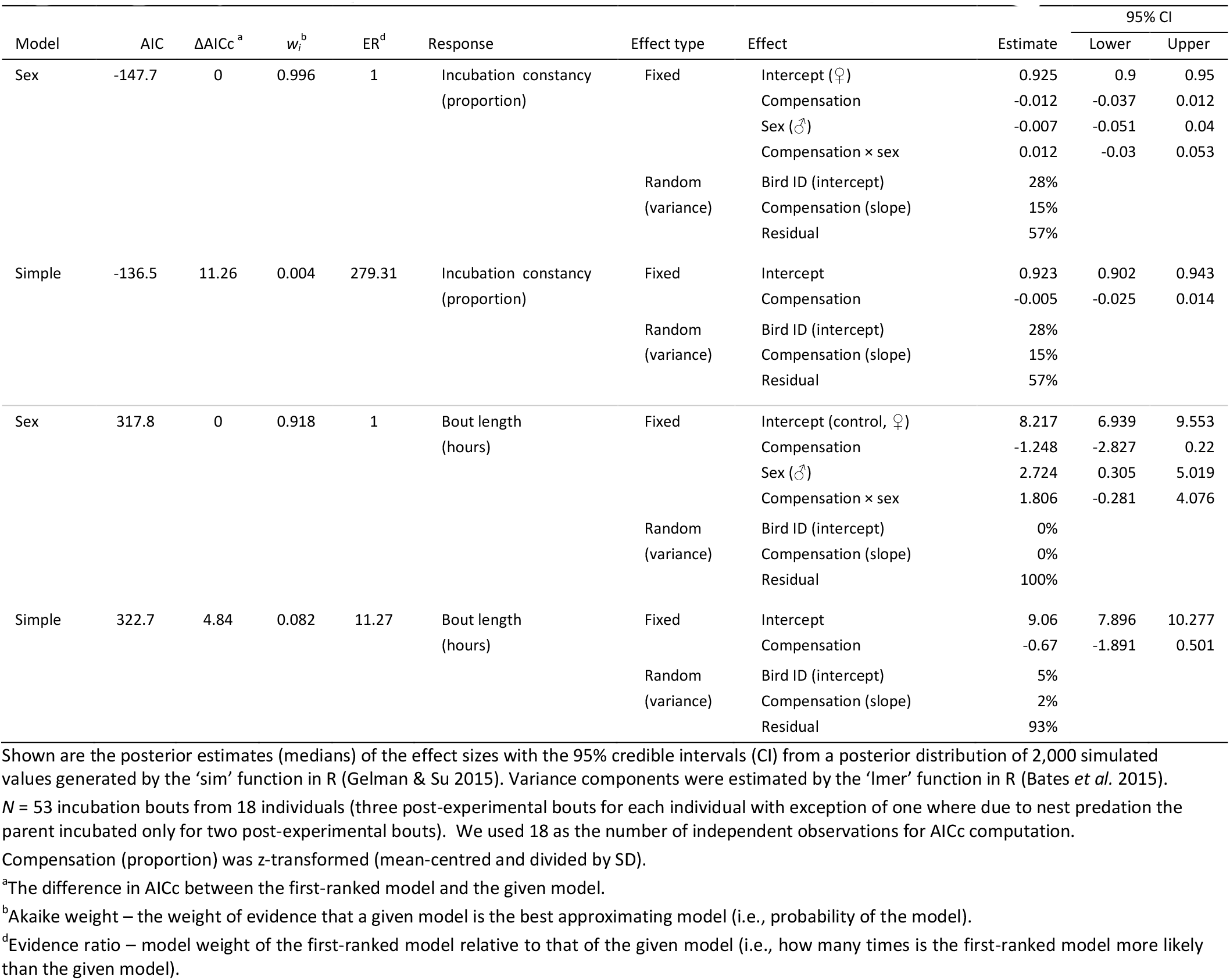
Effect of compensation on focal parent’s nest attendance and bout length in the after-experimental period.

## REFERENCES

Alonso-Alvarez, C. (2001) Effects of testosterone implants on pair behaviour during incubation in the Yellowlegged Gull Larus cachinnans. Journal of Avian Biology, 32, 326–332. http://doi.org/10.1111/¡.0908-8857.2001.320406.x.

AlRashidi, M., Kosztolányi, A., Shobrak, M., Küpper, C. & Székely, T. (2011) Parental cooperation in an extreme hot environment: natural behaviour and experimental evidence. Animal Behaviour, 82, 235–243. http://doi.Org/10.1016/i.anbehav.2011.04.019.

Anderson, D.R. (2008) Model Based Inference in the Life Sciences: A Primer on Evidence. Springer, New York.

Ashkenazie, S. & Safriel, U.N. (1979a) Breeding Cycle and Behavior of the Semipalmated Sandpiper at Barrow, Alaska. The Auk, 96, 56–67. http://doi.org/10.2307/4085399.

Ashkenazie, S. & Safriel, U.N. (1979b) Time-Energy Budget of the Semipalmated Sandpiper Calidris Pusilla at Barrow, Alaska. Ecology, 60, 783–799. http://doi.org/10.2307/1936615.

Bates, D., Maechler, M., Bolker, B. & Walker, S. (2015) Fitting Linear Mixed-Effects Models Using lme4. Journal of Statistical Software, 67, 1–48. http://doi.org/10.18637/iss.v067.i01.

Bell, G.P. (1990) Birds and mammals on an insect diet: a primer on diet composition analysis in relation to ecological energetics. Studies in Avian Biology, 416–422.

Bowman, R. & Bird, D.M. (1987) Behavioral strategies of American kestrels during mate replacement. Behavioral Ecology and Sociobiology, 20, 129–135. http://doi.org/10.1007/bf00572635.

Brunton, D.H. (1988) Sexual differences in reproductive effort: time-activity budgets of monogamous killdeer, Charadrius vociferus. Animal Behaviour, 36, 705–717. http://doi.org/10.1016/s0003-3472(88)80153-2.

Bulla, M. (2014) R-SCRIPT and EXAMPLE DATA to extract incubation from temperature measurements. figshare, https://doi.org/10.6084/m9.figshare.1037545.v1.

Bulla, M. (2017a) Supporting information for 'Flexible parental care: Uniparental incubation in biparentally incubating shorebirds'. Open Science Framework, http://doi.org/10.17605/OSF.IO/3RSNY.

Bulla, M. (2017b) Supporting Information for 'Temporary mate removal during incubation leads to variable compensation in a biparental shorebird'. Open Science Framework, https://osf.io/mx82q/.

Bulla, M., Cresswell, W., Rutten, A.L., Valcu, M. & Kempenaers, B. (2015a) Biparental incubation-scheduling: no experimental evidence for maior energetic constraints. Behavioral Ecology, 26, 30–37. http://doi.org/10.1093/beheco/aru156.

Bulla, M., Prüter, H., Vitnerová, H., Tijsen, W., Sládeček, M., Alves, J.A., Gilg, O. & Kempenaers, B. (2017)Flexible Parental Care: Uniparental Incubation In Biparentally Incubating Shorebirds. bioRxiv. 10.1101/117028.

Bulla, M., Stich, E., Valcu, M. & Kempenaers, B. (2015b) Off-nest behaviour in a biparentally incubating shorebird varies with sex, time of day and weather. Ibis, 157, 575–589. http://dx.doi.org/10.1111/ibi.12276.

Bulla, M., Valcu, M., Dokter, A.M., Dondua, A.G., Kosztolányi, A., Rutten, A., Helm, B., Sandercock, B.K., Casler, B., Ens, B.J., Spiegel, C.S., Hassell, C.J., Küpper, C., Minton, C., Burgas, D., Lank, D.B., Payer, D.C., Loktionov, E.Y., Nol, E., Kwon, E., Smith, F., Gates, H.R., Vitnerová, H., Prüter, H., Johnson, J.A., Clair, J.J.H.S., Lamarre, J.-F., Rausch, J., Reneerkens, J., Conklin, J.R., Burger, J., Liebezeit, J., Bêty, J., Coleman, J.T., Figuerola, J., Hooiimeiier, J.C.E.W., Alves, J.A., Smith, J.A.M., Weidinger, K., Koivula, K., Gosbell, K., Niles, L., Koloski, L., McKinnon, L., Praus, L., Klaassen, M., Giroux, M.-A., Sládeček, M., Boldenow, M.L., Exo, M., Goldstein, M.I., Šálek, M., Senner, N., Rönkä, N., Lecomte, N., Gilg, O., Vincze, O., Johnson, O.W., Smith, P.A., Woodard, P.F., Tomkovich, P.S., Battley, P., Bentzen, R., Lanctot, R.B., Porter, R., Saalfeld, S.T., Freeman, S., Brown, S.C., Yezerinac, S., Székely, T., Montalvo, T., Piersma, T., Loverti, V., Pakanen, V.-M., Tiisen, W. & Kempenaers, B. (2016) Unexpected diversity in socially synchronized rhythms of shorebirds. Nature.

Bulla, M., Valcu, M., Rutten, A.L. & Kempenaers, B. (2013) Data from: Biparental incubation patterns in a high-Arctic breeding shorebird: how do pairs divide their duties? Dryad Data Repository, http://doi.org/10.5061/dryad.nh8f0.

Bulla, M., Valcu, M., Rutten, A.L. & Kempenaers, B. (2014) Biparental incubation patterns in a high-Arctic breeding shorebird: how do pairs divide their duties? Behavioral Ecology, 25, 152–164. http://doi.org/10.1093/beheco/art098.

Burley, N. (1980) Clutch overlap and clutch size: alternative and complementary reproductive tactics. American Naturalist 223–246. http://www.jstor.org/stable/2460595.

Cartar, R.V. & Montgomerie, R.D. (1985) The Influence of Weather on Incubation Scheduling of the White-Rumped Sandpiper (Calidris fuscicollis): A Uniparental Incubator in a Cold Environment. Behaviour, 95, 261–289. http://www.jstor.org/stable/4534487.

Chaurand, T. & Weimerskirch, H. (1994) Incubation routine, body mass regulation and egg neglect in the Blue Petrel Halobaena caerulea. Ibis, 136, 285–290. http://doi.org/10.1111/j.1474-919X.1994.tb01097.x.

Coppens, C.M., de Boer, S.F. & Koolhaas, J.M. (2010) Coping styles and behavioural flexibility: towards underlying mechanisms. Philosophical Transactions of the Royal Society of London B Biological Sciences, 365, 4021–4028. http://doi.org/10.1098/rstb.2010.0217.

De Ridder, E., Pinxten, R. & Eens, M. (2000) Experimental evidence of a testosterone-induced shift from paternal to mating behaviour in a facultatively polygynous songbird. Behavioral Ecology and Sociobiology, 49, 24–30. http://doi.org/10.1007/s002650000266.

Dearborn, D.C. (2001) Body condition and retaliation in the parental effort decisions of incubating great frigatebirds (Fregata minor). Behavioral Ecology, 12, 200–206. http://doi.org/10.1093/beheco/12.2.200.

Deeming, D.C. (2002) Behaviour patterns during incubation. Avian incubation, behaviour, environment and evolution (ed. D.C. Deeming), pp. 63–87. Oxford University Press, Oxford.

Duckworth, J.W. (1992) Effects of mate removal on the behaviour and reproductive success of Reed Warblers Acrocephalus scirpaceus. Ibis, 134, 164–170. http://doi.org/10.1111/j.1474-919X.1992.tb08393.x.

Erckmann, W.J. (1981) The evolution of sex-role reversal and monogamy in shorebirds. PhD PhD, Univ. Washington.

Gauthier-Clerc, M., Le Maho, Y., Gendner, J.-P., Durant, J. & Handrich, Y. (2001) State-dependent decisions in long-term fasting king penguins, Aptenodytes patagonicus, during courtship and incubation. Animal Behaviour, 62, 661–669. http://doi.org/10.1006/anbe.2001.1803.

Gelman, A. & Hill, J. (2007) Data analysis using regression and multilevel/hierarchical models. Cambridge University Press, Cambridge.

Gelman, A. & Su, Y.-S. (2015) arm: Data Analysis Using Regression and Multilevel/Hierarchical Models. R package version 1.8-6., http://CRAN.R-project.org/package=arm.

Gibbon, J., Morrell, M. & Silver, R. (1984) Two kinds of timing in circadian incubation rhythm of ring doves. American Journal of Physiology-Regulatory, Integrative and Comparative Physiology, 247, R1083–R1087.

Harrison, F., Barta, Z., Cuthill, I. & Székely, T. (2009) How is sexual conflict over parental care resolved? A meta-analysis. Journal of Evolutionary Biology, 22, 1800–1812. http://doi.org/10.1111/¡.1420-9101.2009.01792.x.

Hicklin, P. & Gratto-Trevor, C.L. (2010) Semipalmated Sandpiper (Calidris pusilla). The Birds of North America Online. (ed. A. Poole). Cornell Lab of Ornithology, Ithaca.

Honaker, J., King, G. & Blackwell, M. (2011) Amelia II: A Program for Missing Data. Journal of Statistical Software, 45, 1–47.

Houston, A.I. & Davies, N.B. (1985) The evolution of cooperation and life history in the dunnock Prunella modularis. Behavioural ecology: ecological consequences of adaptive behaviour (eds R.M. Sibly & R.H. Smith), pp. 471–487. Blackwell Scientific Publications, Oxford.

Johnstone, R.A. & Hinde, C.A. (2006) Negotiation over offspring care—how should parents respond to each other’s efforts? Behavioral Ecology, 17, 818–827. http://doi.org/10.1093/beheco/arl009.

Jones, K.M., Ruxton, G.D. & Monaghan, P. (2002) Model parents: is full compensation for reduced partner nest attendance compatible with stable biparental care? Behavioral Ecology, 13, 838–843. http://doi.org/10.1093/beheco/13.6.838.

Kersten, M. & Piersma, T. (1986) High levels of energy expenditure in shorebirds: metabolic adaptations to an energetically expensive way of life. Ardea, 75, 175–188. http://dx.doi.org/10.5253/arde.v75.p175.

Koolhaas, J., Korte, S., De Boer, S., Van Der Vegt, B., Van Reenen, C., Hopster, H., De Jong, I., Ruis, M. & Blokhuis, H. (1999) Coping styles in animals: current status in behavior and stress-physiology. Neuroscience & Biobehavioral Reviews, 23, 925–935. http://doi.org/10.1016/S0149-7634(99)00026-3.

Kosztolanyi, A., Cuthill, I.C. & Szekely, T. (2009) Negotiation between parents over care: reversible compensation during incubation. Behavioral Ecology, 20, 446–452. http://doi.org/10.1093/beheco/arn140.

Kosztolányi, A., Székely, T. & Cuthill, I.C. (2003) Why do both parents incubate in the Kentish plover? Ethology, 109, 645–658. http://doi.org/10.1046/j.1439-0310.2003.00906.x.

Lessells, C.M. (2012) Sexual conflict. The evolution of parental care (eds J.A. Royle, P.T. Smiseth & M. Kölliker),pp. 150–170. Oxford University Press, Oxford.

Løfaldli, L. (1985) Incubation rhythm in the great snipe Gallinago media. Ecography, 8, 107–112. http://doi.org/10.1111/¡.1600-0587.1985.tb01160.x.

Mazerolle, M.J. (2016) AICcmodavg: Model selection and multimodel inference based on (Q)AIC(c). R package version 2.0-4.

McDonald, P.G., Buttemer, W.A. & Astheimer, L.B. (2001) The Influence of Testosterone on Territorial Defence and Parental Behavior in Male Free-Living Rufous Whistlers, Pachycephala rufiventris. Hormones and Behavior, 39, 185–194. http://doi.org/10.1006/hbeh.2001.1644.

McNamara, J.M., Gasson, C.E. & Houston, A.I. (1999) Incorporating rules for responding into evolutionary games. Nature, 401, 368–371. http://www.nature.com/nature/iournal/v401/n6751/abs/401368a0.html.

McNamara, J.M., Houston, A.I., Barta, Z. & Osorno, J.-L. (2003) Should young ever be better off with one parent than with two? Behav. Ecol., 14, 301–310. http://doi.org/10.1093/beheco/14.3.301.

Nakagawa, S. & Freckleton, R.P. (2011) Model averaging, missing data and multiple imputation: a case study for behavioural ecology. Behavioral Ecology and Sociobiology, 65, 103–116. http://doi.org/10.1007/s00265-010-1044-7.

Nord, A., Sandell, M.I. & Nilsson, J.-Å. (2010) Female zebra finches compromise clutch temperature in energetically demanding incubation conditions. Functional Ecology, 24, 1031–1036. http://doi.org/10.1111/j.1365-2435.2010.01719.x.

Norton, D. (1973) Ecological Energetics of Calidridine Sandpipers Breeding in Northern Alaska. Doctor of Philosophy, University of Alaska.

Pinxten, R., Eens, M. & Verheyen, R.F. (1995) Response of male European starlings to experimental removal of their mate during different stages of the breeding cycle. Behaviour, 132, 301–317. http://www.jstor.org/stable/4535265.

Poole, A. (2005) The Birds of North America Online. (ed. A. Poole). Cornell Laboratory of Ornithology, Ithaca, NY.

R-Core-Team (2016) R: A Language and Environment for Statistical Computing. Version 3.3.0. R Foundation for Statistical Computing, http://www.R-project.org/.

Reneerkens, J., Grond, K., Schekkerman, H., Tulp, I. & Piersma, T. (2011) Do uniparental Sanderlings Calidris alba increase egg heat input to compensate for low nest attentiveness? PLoS ONE, 6, e16834. http://doi.org/10.1371/journal.pone.0016834.

Schwagmeyer, P.L., Schwabl, H.G. & Mock, D.W. (2005) Dynamics of biparental care in house sparrows: hormonal manipulations of paternal contributions. Animal Behaviour, 69, 481–488. http://doi.org/10.1016/j.anbehav.2004.04.017.

Trivers, R. (1972) Parental investment and sexual selection. Sexual selection and the descent of man (ed. B. Cambell), pp. 136–179. Aldine Press, Chicago.

Vleck, C.M. (1981) Energetic Cost of Incubation in the Zebra Finch. The Condor, 83, 229–237. http://www.jstor.org/stable/1367313.

Weimerskirch, H. (1995) Regulation of foraging trips and incubation routine in male and female wandering albatrosses. Oecologia, 102, 37–43. http://doi.org/10.1007/bf00333308.

Wiebe, K.L. (2010) Negotiation of parental care when the stakes are high: experimental handicapping of one partner during incubation leads to short-term generosity. Journal of Animal Ecology, 79, 63–70. http://doi.org/10.1111/j.1365-2656.2009.01614.x.

Williams, J.B. (1996) Energetics of avian Incubation. Avian Energetics and Nutritional Ecology (ed. C. Carey), pp. 375–416. Chapman & Hall, New York.

## References

Bates, D., Maechler, M., Bolker, B. & Walker, S. (2015) Fitting Linear Mixed-Effects Models Using lme4. Journal of Statistical Software, 67, 1–48. http://doi.org/10.18637/jss.v067.i01.

Bulla, M., Cresswell, W., Rutten, A.L., Valcu, M. & Kempenaers, B. (2015) Biparental incubation-scheduling: no experimental evidence for major energetic constraints. Behavioral Ecology, 26, 30–37. http://doi.org/10.1093/beheco/aru156.

